# Contextual control of CD8^+^ T cell priming by dendritic cell subsets in tumor and inflammatory microenvironments

**DOI:** 10.1101/2025.10.31.685861

**Authors:** Victoria P Schuster, Naomi Berkowitz, Kasidy Brown, Catherine Rousculp, Valentina Laverde, Julia R Unsworth, Megan K Ruhland

## Abstract

Conventional dendritic cells orchestrate adaptive immunity by trafficking peripheral antigens to draining lymph nodes and presenting peptide–MHC complexes to prime naïve T cells. Migratory and lymph node resident conventional dendritic cell subsets occupy distinct anatomical niches and have been shown to shape T cell activation in a variety of immunologic contexts including infection, vaccination and cancer. How peripheral tissue context and dendritic cell subset specific transcriptional programs collaborate to determine CD8^+^ T cell priming remains incompletely defined. Using fluorescent antigen, we tracked antigen distribution, dendritic cell transcriptional programming, and functional cross-presentation across tumor, inflammatory, and steady state tissue contexts. We find that skin tumor antigen is more widely distributed amongst draining lymph node conventional dendritic cells than skin antigen derived from either steady state or inflamed skin tissue. Comparing across dendritic cell subsets, migratory type 1 dendritic cells display higher expression of MHCI antigen presentation machinery and genes associated with cross-presentation compared with lymph node resident type 1 dendritic cells in tumor and inflamed tissue contexts. Similarly, they exhibited superior per-cell cross-presentation and stronger induction of naïve CD8^+^ T cell responses. We find that both antigen access in the lymph node and context-specific cross-presentation efficiency together predict the magnitude and quality of CD8^+^ T cell priming. These results identify migratory dendritic cells, particularly type 1, as central mediators of antitumor CD8^+^ T cell responses and support therapeutic strategies that augment the efficiency of resident dendritic cell cross-presentation.

## INTRODUCTION

Conventional dendritic cells (cDCs) regulate induction of T cell responses through presentation of tissue-derived antigen alongside tolerogenic or activating signals to appropriately activate T cells. Priming of naïve antigen-specific T cells occurs in the lymph nodes (LNs), and migratory cDCs have been shown to traffic exogenous antigen from the peripheral tissue to the LN to drive a productive T cell response^1^. In order to generate CD8^+^ T cell responses to exogenous antigens, migratory cDCs process and present peptide-MHCI complexes on the cell surface through a process termed cross-presentation. Migratory type 1 cDCs (mDC1s) are particularly adept at stimulating CD8^+^ T cell responses through cross-presentation. This function is critical for antiviral responses, antitumor immunity, and maintenance of self tolerance^1–3^. While migratory type 2 cDCs (mDC2s) are generally considered to be more efficient at driving CD4^+^ T cell responses, they have also been shown to cross-present tumor antigen to CD8^+^ T cells^4–6^. However, in the context of solid tumors, mDC1s have been shown to be indispensable for stimulating antitumor CD8^+^ T cell responses across many murine tumor models^6,7^. Importantly, this is also reflected in human biology, where high mDC1 signatures, not mDC2 signatures, correlate with improved CD8^+^T cell tumor infiltration and patient survival^3,8^.

Following the transport of antigen to the LN, migratory cDC subsets are capable of transferring antigen as discrete vesicular packages to LN resident type 1 (rDC1) or type 2 (rDC2) cDCs^9,10^. During this transfer of antigen from migratory to resident cDCs, contextual cues derived from the tissue can also be passed, including co-transfer of receptors within the vesicular package^11^. This additional information regarding tissue context can facilitate synchronous T cell priming by both LN resident and migratory cDCs. However, understanding of the patterns and degree of antigen transfer from migratory to resident cDCs in different inflammatory contexts remains limited. In model systems where transfer has been well characterized, it is clear that the state of peripheral tissues heavily influences the dynamics of antigen passing in the draining LN (dLN)^12^. These findings suggest that resident cDCs play a differential role in instructing the immune response that is dependent on the state of the peripheral tissue. Studies in viral immunity suggest that antigen transfer to rDC1s is essential for immunity^9,13^. However, in antitumor immunity, studies suggest that rDC1s can drive suboptimal CD8^+^ T cell responses^10^.

In any context, two main factors contribute to the ability of a given cDC to prime a naïve CD8^+^ T cell: 1) access to antigen, and 2) efficiency of cross-presentation. Understanding both of these aspects of cDC biology will be essential for designing therapeutic strategies that optimize T cell priming. Here we used model systems that enable antigen tracking to determine how antigen transfer to dLN resident cDCs contributes to CD8^+^ T cell priming in different skin tissue contexts and to evaluate the efficiency and context-specific, cell-intrinsic differences in cross-presentation across migratory and resident cDC subsets. While previous work has studied cDC subsets within individual models, here we compare CD8^+^ T cell activation across tumor, inflammation and steady state to identify characteristics that may enable context-specific, therapeutic targeting of cDCs.

## RESULTS

### Skin tumor antigen is more dispersed than antigen from steady state or inflamed skin in dLN cDCs

To evaluate the dynamics of antigen dispersal in dLN cDC subsets across distinct peripheral tissue contexts, we used K14-Cre; lox-stop-lox-ZsGreen mice (K14-ZsGreen) at steady state, K14-ZsGreen mice treated topically with the inflammatory inducer dibutyl phthalate (DBP), and wildtype C57BL/6 mice subcutaneously implanted with B16F10 melanoma cells expressing cytoplasmic ZsGreen (B16-ZsGreen). DBP is a commonly used murine skin sensitizing agent that facilitates the activation of Th2 inflammatory responses^12,14^. These models enabled tracking of skin-derived fluorescent antigen from keratinocytes (K14-ZsGreen) or tumor (B16-ZsGreen) cells (Fig. 1A)^6,10^. Skin dLNs (inguinal, axillary and brachial) were harvested from steady state and DBP-treated mice after 7 days of treatment, and tumor-bearing mice 14 days after implantation. Skin dLNs were digested and stained with an antibody panel to identify cDC subsets and assess antigen acquisition (i.e. ZsGreen^+^) by flow cytometry (Fig. 1A, Fig. S1A).

**Figure 1.**
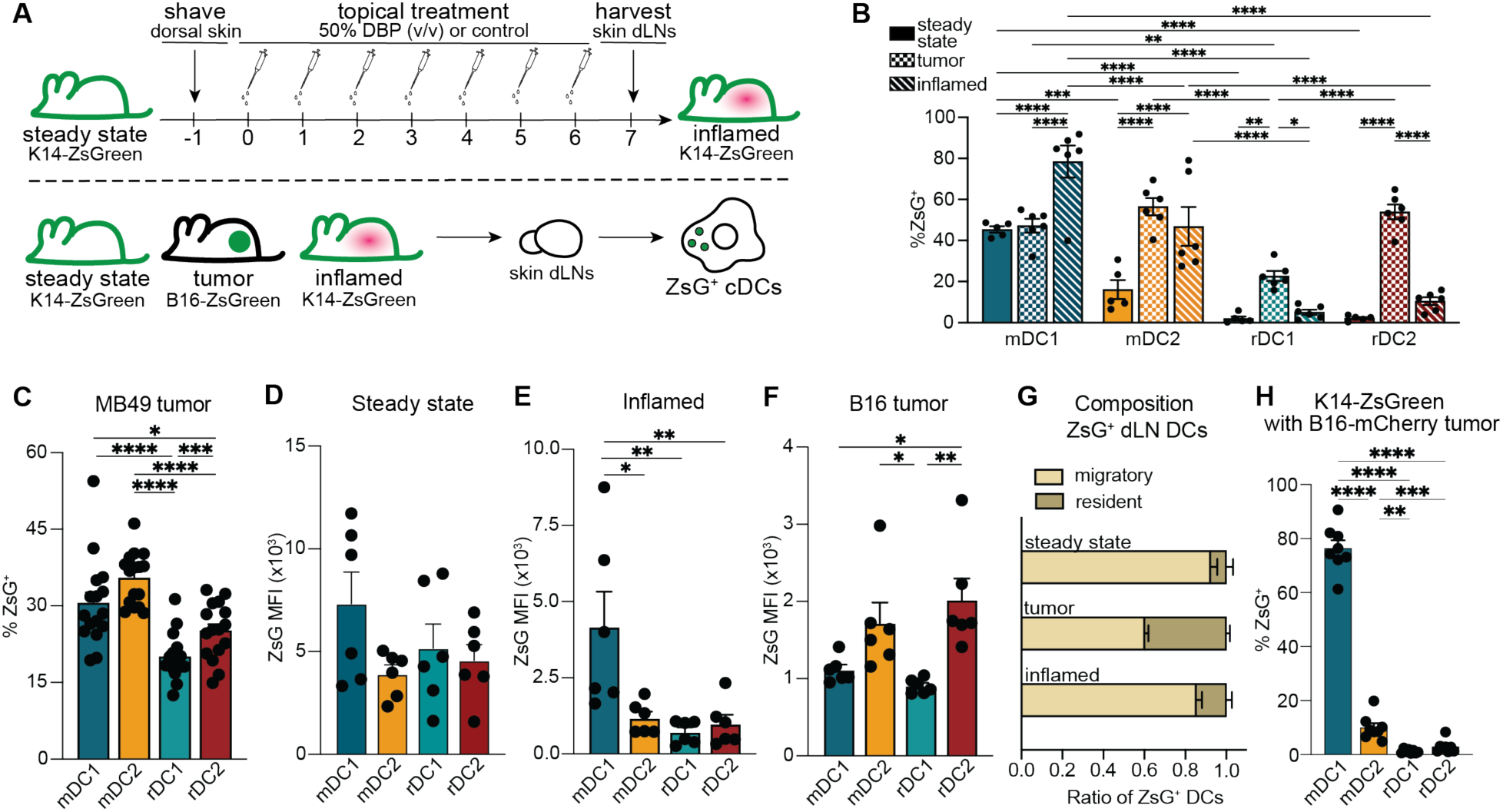
Peripheral tissue state determines antigen dispersal amongst cDC subsets in dLNs. (**A**) Schematic for (top) topical application of 50% (v/v) dibutyl phthalate (DBP) or vehicle (acetone) control to induce inflamed skin model or maintain steady state skin and (bottom) flow cytometry analysis of conventional dendritic cell (cDC) subsets from draining lymph nodes (dLNs) of steady state K14-cre; lox-stop-lox-ZsGreen (K14-ZsGreen), inflamed K14-ZsGreen, or C57Bl/6 mice bearing subcutaneous B16-F10 tumors expressing ZsGreen (B16-ZsGreen). (**B - C**) Flow cytometry analysis for percent ZsGreen^+^ cDC composition from dLN cDC subsets of (B) steady state, B16-ZsGreen tumor, or inflamed tissues or (C) MB49-ZsGreen tumors. (**D - F**) Mean fluorescence intensity (MFI) of ZsGreen^+^ cDC populations from dLNs of (D) steady state, (E) inflamed, and (F) B16-ZsGreen tumor bearing mice. (**G**) Ratio of ZsGreen^+^ migratory or resident cDC subsets harvested from dLNs of steady state, B16-ZsGreen tumor and inflamed skin tissue. (**H**) Percent ZsGreen^+^ cDC subsets from dLNs of K14-ZsGreen mice implanted with B16-mCherry tumor on day 14 post-implantation. (B - H) Data are plotted as mean ± SEM. (B, D - G) n = 5 (steady state) or n = 6 (inflamed or B16-ZsGreen tumor) mice (6 lymph nodes each) per tissue condition, data pooled from 2 independent experiments. (C, H) Draining lymph nodes of (C) n = 16 or (H) n = 8 tumors, data pooled from 2 independent experiments. *p < 0.05, **p < 0.01, ***p < 0.001, ****p < 0.0001 by (B) ordinary two-way ANOVA, (C, H) RM one-way ANOVA with Geisser-Greenhouse correction, or (D - F) ordinary one-way ANOVA with (B - F, H) Tukey’s multiple comparisons test. Migratory type 1 cDC, mDC1. Migratory type 2 cDC, mDC2. Resident type 1 cDC, rDC1. Resident type 2 cDC, rDC2. ZsGreen, ZsG.

The relative abundance of individual cDC subsets was largely consistent across conditions, with only a modest decrease in the rDC2 population in tumor-bearing mice compared with inflamed mice (Fig. S1B). In contrast, the frequency of antigen-bearing cDCs differed significantly across peripheral tissue contexts (Fig. 1B). At steady state, as shown previously^10^, the mDC1 subset had the highest percentage of antigen^+^ cells, with approximately 50% of mDC1s containing detectable antigen compared with fewer than 20% of mDC2s, while resident cDC populations exhibited negligible antigen positivity (Fig. 1B). Consistent with ongoing sampling of peripheral tissues by migratory cDCs, these data indicate that steady state antigen remains largely restricted to migratory populations with minimal transfer to resident cDCs. Interestingly, DBP-inflamed skin dLNs led to significantly increased antigen^+^ migratory cDC1s and cDC2s within dLNs compared to steady state, while maintaining limited transfer of antigen to resident cDC populations (Fig. 1B). A minimal increase of antigen^+^ resident cDCs was observed in the inflamed condition compared to steady state (Fig. S1C). However, this increase did not reach statistical significance for the individual resident cDC subsets (Fig. 1B). Analysis of DBP-treated mice at an early inflammatory timepoint (day 3) and a late timepoint (day 10) revealed persistently limited antigen accumulation within resident cDC subsets (Fig. S1D). Thus, despite increases in antigen^+^ migratory cDCs, inflamed conditions did not promote substantial antigen transfer to resident cDC populations throughout the course of the response.

By comparison, tumor dLNs displayed broad distribution of antigen amongst cDC subsets. Across tumor dLN cDC subsets, we observed similar frequencies of antigen^+^ mDC1s, mDC2s and rDC2s; only rDC1s showed a significantly lower percentage of tumor antigen loading than the migratory cDC subsets (Fig. 1B). Resident cDC populations contained significantly higher proportions of antigen-loaded cells than those observed in either steady state or inflamed conditions (Fig. 1B, S1E). Importantly, ZsGreen signal was concentrated within CD45^+^ cells within the tumor dLN and was largely absent from CD45^-^ cells, arguing against substantial passive drainage of tumor-derived fluorescent material or disseminated tumor cells (Fig. S1F). This was consistent with our previous intravital and confocal imaging studies, which demonstrated intracellular localization of ZsGreen and CCR7-dependent delivery of tumor antigen to dLNs^10^. Together, these data support the interpretation that tumor-derived antigen is actively trafficked to the dLN and dispersed among cDC subsets.

To determine whether this antigen dispersal pattern was dependent on the amount of tumor burden, we evaluated earlier tumor-bearing timepoints. Broad distribution of antigen across cDC subsets was apparent on day 8 post-implantation when tumors were small (Fig S1G) and remained largely unchanged at days 10 and 14, indicating that antigen dispersal is established early during tumor development and persists as tumors progress (Fig. S1H). Importantly, a similar pattern of antigen distribution was observed in mice bearing ZsGreen-expressing MB49 tumors (MB49-ZsGreen), demonstrating that widespread antigen dispersal among cDC subsets is not unique to the B16F10 melanoma model (Fig. 1C).

Analysis of antigen quantity within antigen^+^ cells further highlighted differences across tissue contexts. At steady state, all antigen-bearing cDC subsets contained comparable amounts of antigen based on mean fluorescence intensity (MFI) of ZsGreen, despite differences in the frequency of antigen^+^ cells (Fig. 1D). Inflamed conditions increased the relative antigen load within mDC1s, which contained significantly greater amounts of antigen than other subsets (Fig. 1E), consistent with the established role of migratory cDC1s in transporting peripheral antigen to dLNs during inflammation^9,15^. In contrast, tumor dLNs showed a distinct pattern in which antigen levels varied across antigen^+^ subsets (Fig. 1F). Consistent with this, a larger proportion of resident cDC populations contained antigen in the tumor dLN than in either steady state or inflamed conditions (Fig. 1G). Together, these findings reveal a shift in antigen distribution across cDC subsets depending on the peripheral tissue context. Whereas steady state antigen remains largely restricted to migratory cDCs, inflammation increases antigen acquisition by these migratory populations without substantial transfer to resident cDCs. Tumor dLNs display extensive antigen dispersal throughout cDC subsets, resulting in increased antigen within resident populations and a more uniform distribution of antigen among cDC subsets.

We next asked whether tumors generally alter antigen handling within the dLNs or whether enhanced dispersal is specific to tumor-derived antigen. To address this question, B16F10 tumors expressing the fluorescent protein mCherry were implanted into K14-ZsGreen mice, enabling assessment of keratinocyte-derived antigen during tumor growth. Despite the altered peripheral environment, skin-derived antigen remained preferentially restricted to migratory cDC populations, with little accumulation in resident cDCs (Fig. 1H). Thus, the presence of a tumor is not sufficient to enhance transfer of all antigen types within the dLN. Instead, the extensive dispersal observed in tumor-bearing mice appears to be selective for tumor-derived antigen, further supporting a model in which tumor antigens are actively trafficked from the periphery and subsequently redistributed among dLN cDC populations.

Collectively, these data demonstrate that both inflammation and tumors increase antigen acquisition by migratory cDCs; however, robust transfer of antigen to resident cDC populations occurs specifically in the context of tumors. These data suggest that antigen dispersal within tumor dLNs is distinct from that observed during peripheral inflammation, which may impact cross-presentation and characteristics of T cell priming.

### Tissue environment and cDC subtype both contribute to CD8^+^ T cell priming

DLNs displayed different antigen dispersal patterns amongst cDC subsets based on peripheral environment (Fig. 1B, 1G). Given the critical role of antigen availability for cDC-mediated CD8^+^ T cell priming, we next sought to determine whether differential antigen dispersal to resident cDCs impacted CD8^+^ T cell priming characteristics. To track antigen-specific CD8^+^ T cell priming while still allowing *in vivo* cDC antigen loading and dispersal, we used the model antigen ovalbumin (OVA). Specifically, we generated a K14-cre; lox-stop-lox-ZsGreen; K14-OVA mouse model where K14-expressing keratinocytes express both ZsGreen and OVA to evaluate T cell priming against skin-derived antigen. We leveraged the B16-ZsGreen-OVA tumor cell line to study tumor antigen-specific T cell priming^6^. We treated mice with either vehicle (steady state) or DBP (inflamed) or implanted B16-ZsGreen-OVA tumors. ZsGreen^+^ cDC subsets from each condition were isolated via FACS and cocultured with proliferation dye-labeled, OVA-specific, OTI CD8^+^ T cells (Fig. 2A, S2A). Following 72 hours of coculture, we next sought to determine whether the proliferating T cells were functionally different based on the subtype and peripheral tissue conditions of the cross-presenting cDC. Due to the very low numbers of ZsGreen^+^ resident cDCs at steady state (Fig. 1B), it was not possible to obtain sufficient numbers of resident cDCs for T cell stimulation assays. To determine how priming of naïve CD8^+^ T cells by cDC subsets from different tissue contexts impacts the T cells, we used surface and intracellular antibody staining to evaluate T cell functional markers. Expression and nuclear localization of transcription factors T-bet and Eomes augment CD8^+^ T cell effector function, in part through their competitive, repressive binding within the promoter of *Pdcd1*, a gene encoding PD-1^16–18^. At steady state, priming of naïve CD8^+^ T cells by mDC1s led to lower levels of both Eomes and PD-1 expression compared to mDC2 priming (Fig. S2B-C). Within the inflamed condition, mDC1 priming also drove decreased PD-1 and Eomes expression in proliferating CD8^+^ T cells compared to T cells primed by resident cDCs (Fig. S2B-C). In tumor conditions, the increase in Eomes, a weak repressor of PD-1, was not observed. However, an increase in T-bet expression coincided with increased PD-1 expression on CD8^+^ T cells primed by tumor-draining rDC1s compared to mDC1s (Fig. S2D). Combined, these data may suggest a minor improvement in differentiation of T cell effector populations from priming by mDC1s^16–18^. However, this difference was lost when comparing across the subsets and tissue contexts, indicating context dependent differences between subset priming may ultimately be minimal (Fig. S2E). Beyond differentiation, we also sought to determine functional differences of CD8^+^ T cells by assessing expression levels of effector cytokines IFNγ, TNFα, and Granzyme B. Both at steady state and in inflamed conditions, proliferating CD8^+^ T cell production of effector molecules was not affected by cDC subsets (Fig. S2F-H). While an increase in the the percentage of IFNγ^+^ T cells was observed when T cells were primed by tumor dLN derived rDC1s compared to mDC1s (Fig. S2F), this did not correspond with an increase in other effector molecules (Fig. S2G-H). Furthermore, when comparing across subsets and tissues, there were no significant differences between the effector cytokine profiles of the proliferating CD8^+^ T cells (Fig. S2I).

**Figure 2.**
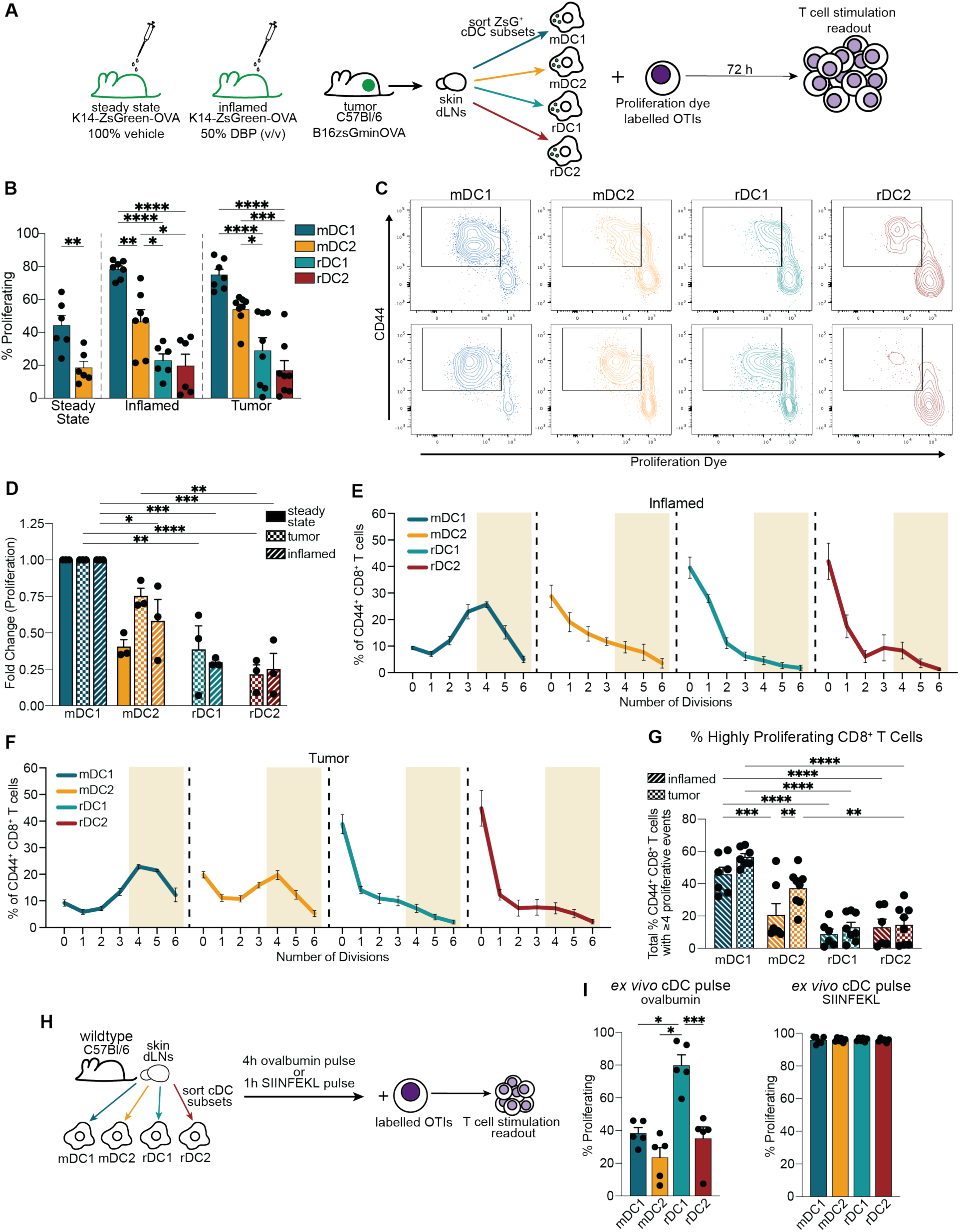
mDC subsets drive productive CD8^+^ T cell proliferation in steady state, inflamed and tumor contexts. (**A**) Schematic for proliferation assay under different tissue conditions. (**B**) Percent proliferating OTI CD8^+^ T cells following coculture with ZsGreen^+^ cDC subsets sorted from skin dLNs of mice under steady state K14-ZsGreen-OVA, inflamed K14-ZsGreen-OVA, or B16-ZsGreen-OVA tumor conditions as shown in A. (**C**) Representative flow cytometry plots depicting T cell proliferation for (top) inflamed and (bottom) tumor conditions following coculture with ZsGreen^+^ mDC1, mDC2, rDC1, or rDC2 subsets. (**D**) Percent proliferating OTIs following ZsGreen^+^ cDC cocultures normalized to fold change over mean mDC1-induced proliferation in a given experiment. (**E - G**) Proliferative modeling by cDC subsets from (E) inflamed or (F) tumor conditions, and (G) Percent highly proliferative CD8^+^ T cells across tissue conditions. Highly proliferative as defined by 4 proliferative events, indicated by shaded regions in (E) and (F). (**H**) Schematic for sorted steady state cDC subset incubations with whole ovalbumin protein or peptide (SIINFEKL) followed by OTI T cell proliferation assay. (**I**) Percent OTI T cell proliferation following coculture with sorted cDC subsets incubated *ex vivo* with (left) whole ovalbumin or (right) SIINFEKL pulsed cDC subsets from steady state mice. (B, D - G, I) Data are plotted as mean ± SEM. (B, D - G) One to three technical replicates from cDCs sorted from 10 mice (6 lymph nodes each) per tissue condition for each experiment, n = 6 - 8 from pooled data from 3 independent experiments. (I) Two to three technical replicates from cDCs sorted from 15 - 20 mice (6 lymph nodes each) per tissue condition for each experiment, n = 5 from pooled data from 2 independent experiments. *p < 0.05, **p < 0.01, ***p < 0.001, ****p < 0.0001 by (B) individual ordinary one-way ANOVAs, (D, G) ordinary two-way ANOVA, or (I) RM one-way ANOVA with Geisser-Greenhouse correction with (B, D, G, I) Tukey’s multiple comparisons test. Migratory type 1 cDC, mDC1. Migratory type 2 cDC, mDC2. Resident type 1 cDC, rDC1. Resident type 2 cDC, rDC2. ZsGreen, ZsG.

While effector cytokine and transcription factor expression did not appear to differentiate cDC subset primed T cells, proliferation rates showed distinct patterns. Following 72 hours of coculture, we found that steady state antigen^+^ mDC1s drove only a modest level of OTI CD8^+^ T cell proliferation though significantly more than antigen^+^ mDC2s (Fig. 2B, Fig. S2J). Next, we evaluated T cell proliferation driven by cDCs isolated from the inflamed condition. We found that antigen-loaded migratory cDCs drove greater OTI CD8^+^ T cell proliferation than steady state migratory cDCs, with mDC1s achieving the most potent T cell priming as measured by T cell proliferation (Fig. 2B-C). While rDC1s and rDC2s were capable of driving CD8^+^ T cell proliferation when isolated from the inflamed context, this was at a significantly lower level than either mDC1s or mDC2s. In agreement with previous observations^6,10^, tumor antigen-loaded migratory cDCs drove potent OTI T cell proliferation (Fig. 2B-C). Resident cDC subsets from tumor dLNs were also capable of driving CD8^+^ T cell proliferation, but similar to the inflamed condition, proliferation was significantly less robust. Interestingly, there was no significant difference between the priming efficiencies of a given cDC subset across inflamed and tumor conditions (Fig. 2D). This suggests that maturation signals from both the inflamed and tumor environment are sufficient to induce antigen presentation on MHCI regardless of antigen origin. When comparing within conventional subsets (i.e. mDC1 vs rDC1) we found that the migratory subset consistently drove the highest level of cumulative OTI T cell proliferation in both inflamed and tumor contexts (Fig. 2B). In both contexts, CD8^+^ T cells primed by resident cDC subsets underwent a reduced number of proliferative events compared to migratory cDC counterparts (Fig. S2K). Proliferative modeling to analyze the number of induced cell divisions from activated CD8^+^ T cells indicated mDC1s are the most robust at priming CD8^+^ T cells as shown by their ability to induce the greatest number of cell divisions (Fig. 2E-F). Under inflamed conditions, few activated CD44^+^ T cells remained undivided and most had undergone ∼4 divisions when cocultured with mDC1s (Fig. 2E, 2G). However, in mDC2 and resident cDC cocultures, the majority of activated T cells had undergone 0-1 divisions within the same timeframe. Similarly, mDC1s derived from tumor conditions drive ∼4 divisions of naïve OTI T cells within 72 hours (Fig. 2F-G). In contrast, although some CD8^+^ T cells did not proliferate in coculture with mDC2s isolated from tumor dLNs, those that did proliferate underwent a similar number of divisions as those cocultured with mDC1s from tumor dLNs (Fig. 2E-G). Regardless of the context from which the cDCs were isolated (inflamed versus tumor), when OTIs were primed by rDC1s or rDC2s, most failed to be highly proliferative (Fig. 2F-G). Taken together, these data suggest that the cross-presenting cDC subset primarily impacts the kinetics of CD8^+^ T cell proliferation, with less impact on effector phenotype.

While antigen-loaded resident cDCs consistently induced less robust proliferation than their migratory counterparts in both inflamed and tumor contexts, these experiments could not distinguish whether this reflected an intrinsic deficiency in cross-presentation or reduced access to antigen in a form capable of being presented on MHCI. To directly address this question, cDC subsets were isolated from skin dLNs of wildtype mice and cultured *ex vivo* with either whole OVA protein or the OVA peptide (SIINFEKL) prior to coculture with proliferation dye-labeled OTI CD8^+^ T cells (Fig. 2H). When pulsed with SIINFEKL peptide, all cDC subsets induced robust OTI proliferation, confirming that resident and migratory cDC subsets are similarly capable of presenting cognate peptide (Fig. 2I). When provided whole OVA protein, all cDC subsets were capable of cross-presenting antigen and inducing T-cell proliferation to varying degrees; strikingly, rDC1s drove significantly greater proliferation than all other cDC subsets (Fig. 2I). These findings demonstrate that resident cDCs, particularly rDC1s, do not possess an intrinsic defect in antigen cross-presentation. Rather, the reduced priming observed following isolation of antigen-loaded resident cDCs from inflamed and tumor dLNs likely reflects differences in maturation state or the form of antigen acquired through antigen dispersal *in vivo*.

### mDC antigen uptake from tumor leads to preferential increased expression of antigen presentation proteins and cytokines

Resident cDCs from dLNs in either inflamed or tumor contexts were less efficient at priming naïve CD8^+^ T cells *in vitro* than migratory cDC populations (Fig. 2B-G). To discern transcriptional differences within cDC subsets between different tissue contexts, we performed bulk RNA sequencing from sorted dLN cDC subsets that were positive for peripheral antigen (ZsGreen^+^) from either steady state, inflamed or tumor tissue (Fig. 3A). We compared expression of genes known to promote productive T cell proliferation, including genes involved in cross-presentation of cell-derived antigen, including costimulation and cytokine production. Notably, we found expression of key inflammatory cytokines *Il1b* and *IL12b* were signficantly upregulated specifically in mDC1s (p-values of 0.049 and 0.0015, respectively), but not rDC1s, from tumor dLNs (Fig. 3B-C). We sought to validate these findings at the protein level by flow cytometry staining and found that while IL12 protein levels were significantly increased in antigen^+^ mDC1s, but not rDC1s, from tumor dLNs compared to inflamed (Fig. S3A-B), IL1β protein levels were not significantly different (Fig. S3C-D). Production of these cytokines, particularly IL12p70, produced by cDCs have been shown to be key for inducing Th1 polarization and are important for antitumor immunity^19–21^. Interestingly, flow cytometry staining of tumor dLNs showed that the percentage of IL12 expressing cDCs was highest within the mDC1 subset (Fig. S3E). Beyond cytokine differences, we found that antigen-loaded migratory cDC subsets isolated from tumor dLNs had significantly increased expression of multiple MHC genes compared to antigen-loaded migratory cDCs isolated from inflamed contexts (Fig. 3D, 3E). The same MHC transcripts in resident cDCs were not found to be upregulated in tumor dLNs compared to the corresponding subsets in inflamed dLNs (Fig. 3F, 3G).

**Figure 3.**
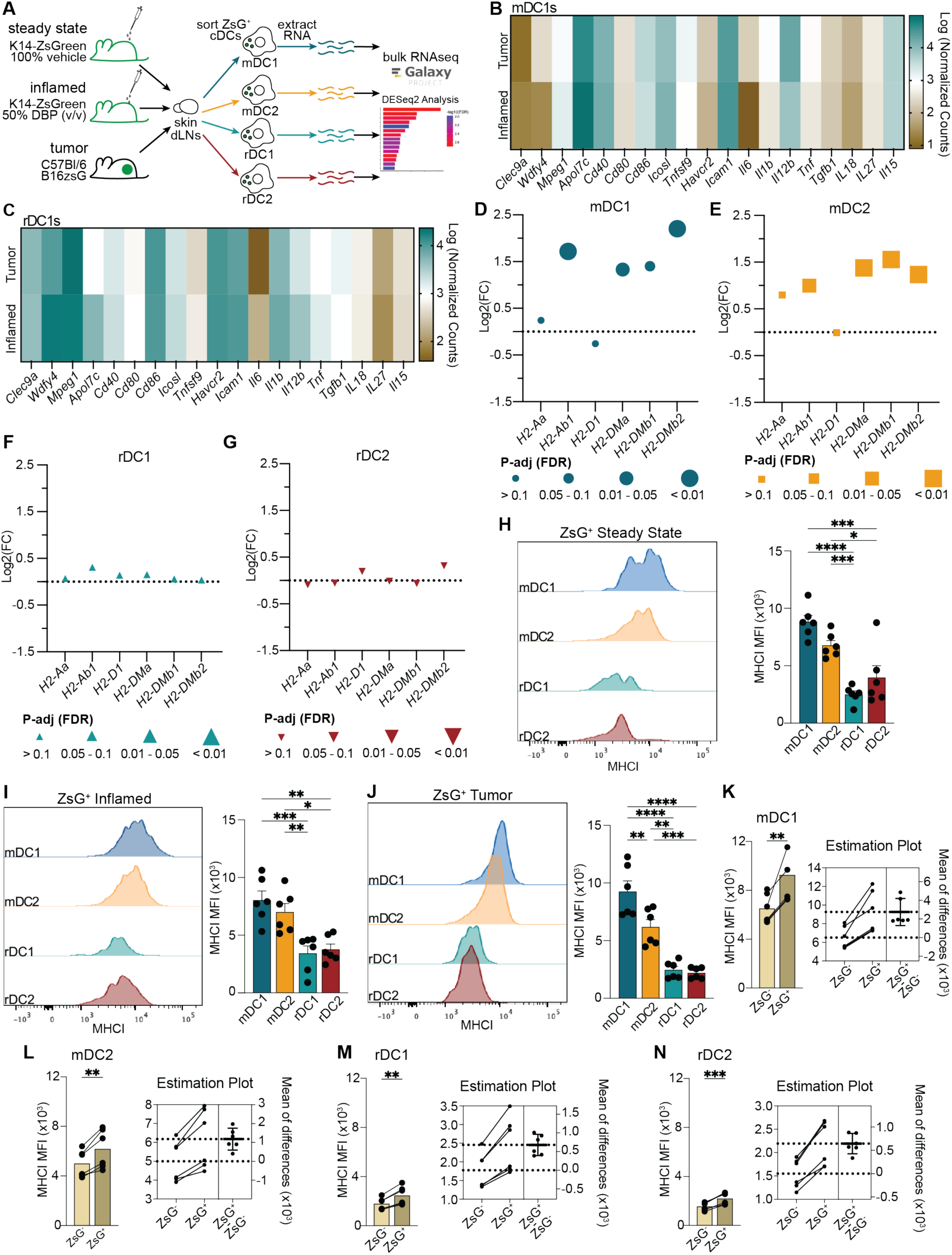
mDCs more effectively upregulate antigen presentation components in tumor conditions. (**A**) Schematic describing bulk RNA sequencing of ZsGreen^+^ cDC subsets sorted from dLNs from steady state, inflamed, or B16-ZsGreen tumor conditions. (**B - C**) Heatmap of normalized counts for gene expression from ZsGreen^+^ (B) mDC1s or (C) rDC1s derived from tumor versus inflamed conditions. (**D - G**) Differentially expressed genes from tumor vs inflamed conditions for antigen presentation genes of sorted ZsGreen^+^ (D) mDC1s, (E) mDC2s, (F) rDC1s, (G) rDC2s from dLNs. (**H - J**) Representative histograms (left) and MFI (right) of MHCI from flow cytometry analysis of ZsGreen^+^ cDC subsets from (H) steady state, (I) inflamed, and (J) tumor conditions. (**K - N**) MHCI (left) MFI and (right) estimation plot of ZsGreen^-^ and ZsGreen^+^ paired subsets of (K) mDC1s, (L) mDC2s, (M) rDC1s, (N) rDC2s from dLNs of tumor bearing mice. (H - J) Data are plotted as mean ± SEM, (B - C) mean, or (K - N) mean ± 95% CI. (B - G) n = 2 technical replicates; each replicate represents 20 mice (6 lymph nodes each) per tissue condition. (H - N) n = 6 mice (6 lymph nodes each) per tissue condition, data pooled from 2 independent experiments. *p < 0.05, **p < 0.01, ***p < 0.001, ****p < 0.0001 by (H - J) ordinary one-way ANOVA with Tukey’s multiple comparisons test or (K - N) Gaussian paired two-tailed t test. Migratory type 2 cDC, mDC2. Resident type 1 cDC, rDC1. Resident type 2 cDC, rDC2. ZsGreen, ZsG.

Previous work highlighted the importance of antigen transfer in vaccine models, suggesting that sharing antigen across cDC populations in the LN boosts antigen availability and drives increased T cell priming^9,22^. The contribution of antigen passing to antitumor T cell priming in the tumor dLN remains unclear^10,11^. When comparing across model systems, we found that antigen-bearing migratory cDCs have significantly higher MHCI surface levels compared to resident cDC subsets, across tissue contexts (Fig. 3H-J). Similar differences were found when evaluating migratory vs resident cDC subset expression of the costimulatory molecule CD86, with migratory cDCs expressing significantly greater levels of CD86 compared to resident populations from tumor dLNs (Fig. S3F, S3G). Furthermore, across conditions, mDCs represented the greatest percentages of CD86^+^ cDCs within the dLNs. Increased MHCI and CD86 in mDCs compared to rDCs in tumor dLNs was also identified with MB49-ZsGreen tumors, suggesting these phenotypes are not restricted to the B16F10 tumor model (Fig. S3H-I). These findings supported the data showing that mDCs drove the greatest proliferation in CD8^+^ T cells *in vitro* (Fig. 2D, 2G). Notably, in tumor conditions, surface level MHCI on mDC1s was significantly higher than that on mDC2 populations, consistent with previous work suggesting mDC1s are crucial for CD8^+^ T cell priming in the context of tumor (Fig. 3J)^6,8,23^. We next asked how the uptake of antigen impacted surface MHCI levels on a per subset basis within the tumor dLN. We found that mDC1s have the highest increase in surface MHCI following antigen uptake (Fig. 3K), at over 40%, compared to mDC1s without tumor antigen. This increase was over 2-fold more than the mDC2 increase (Fig. 3L), and over 4-fold more than the increase seen for either resident cDC population (Fig. 3M, 3N). This subset-specific difference in increased surface MHCI levels upon antigen uptake further supports the specialization of mDC1s for driving optimal antitumor CD8^+^ T cell priming.

Given the superior capacity of rDC1s to cross-present exogenous antigen when provided OVA protein *ex vivo* (Fig. 2H-I), we next sought to determine whether resident cDCs were capable of upregulating antigen presentation and costimulatory molecules in response to *ex vivo* inflammatory stimulation. To test this, we isolated skin-draining LNs from wildtype mice, digested them to obtain single cell suspesions and stimulated the cells with LPS for 4 hours prior to evaluation of MHCI and CD86 expression (Fig. S3J-L). Consistent with the expression patterns observed in tumor dLNs, mDC1s exhibited a significant, albeit modest, increase in surface MHCI following TLR stimulation, whereas no significant increase was observed in mDC2s, rDC1s, or rDC2s (Fig. S3K). In contrast, LPS induced only a small increase in CD86 expression in rDC1s, with little to no change observed in the remaining cDC subsets (Fig. S3L). These data indicate that resident cDCs are not intrinsically incapable of antigen presentation, as demonstrated by their ability to efficiently cross-present OVA protein (Fig. 2H-I). Rather, resident cDCs may be comparatively limited in their ability to upregulate MHCI-associated antigen presentation programs in response to tumor antigen uptake and/or inflammatory stimulation.

Taken together, these data show that antigen uptake is associated with subset specific increases in MHC gene expression and cytokine expression specific to mDC1s within the context of tumor (Fig 3B, 3D). In contrast, rDC1 expression of MHCI remains constant between tumor and inflamed tissue states (Fig. 3C, 3F). Further, migratory cDC populations, specifically mDC1s, are particularly well-suited for presenting antigen from tumor environments compared to resident populations due to high expression of MHCI, IL12 and CD86 (Fig. 3J, S3E-F).

### MHCI machinery and associated molecular pathways are upregulated in mDC1s vs rDC1s in tissues

Type 1 cDCs are strongly implicated in CD8^+^ T cell responses in both inflamed and tumor conditions^23–25^. Our intra-subset comparison of cDC1s between tissue contexts revealed rDC1s have less variable expression of cross-presentation genes between tumor and inflamed tissue states compared to the increased expression of those genes in tumor dLN mDC1s. Given evolving insights into migratory and resident cDC1 subsets and their progenitor populations^26–28^, parsing out subset intrinsic gene expression differences across different tissue contexts may help refine their relative contributions to initiating crucial CD8^+^ T cell responses. Using RNA sequencing, we compared the mDC1s versus rDC1s within individual tissue states. We first compared mDC1s to rDC1s in an inflamed tissue response (Fig. 4A). ShinyGo (v.0.82) was used to determine which cellular components were being upregulated in mDC1s. Compared to rDC1s, mDC1s primarily upregulated expression of the MHCI protein complex and associated proteins (Fig. 4B). mDC1s also had higher expression of the CD40 costimulatory molecule complex. Both of these components are important for T cell activation and antigen-specific immune responses. This was also reflected in the top biological pathways upregulated in mDC1s versus rDC1s from inflamed conditions. Specifically, the pathways involved in peptide transport and formation of peptide-MHCI and peptide-MHCI-TCR bound complexes were upregulated in mDC1s (Fig. 4C). Further analysis of the cellular components involved in the peptide-MHCI loading complex showed specific gene upregulation primarily of the MHCI molecules in mDC1s and much of the associated machinery (Fig. 4D). Together these upregulated cellular components, pathways and MHCI genes suggest mDC1s may have increased interactions with CD4^+^ T cells via CD40 and CD8^+^ T cells through MHCI binding, likely contributing to improved DC licensing and CD8^+^ T cell priming.

**Figure 4.**
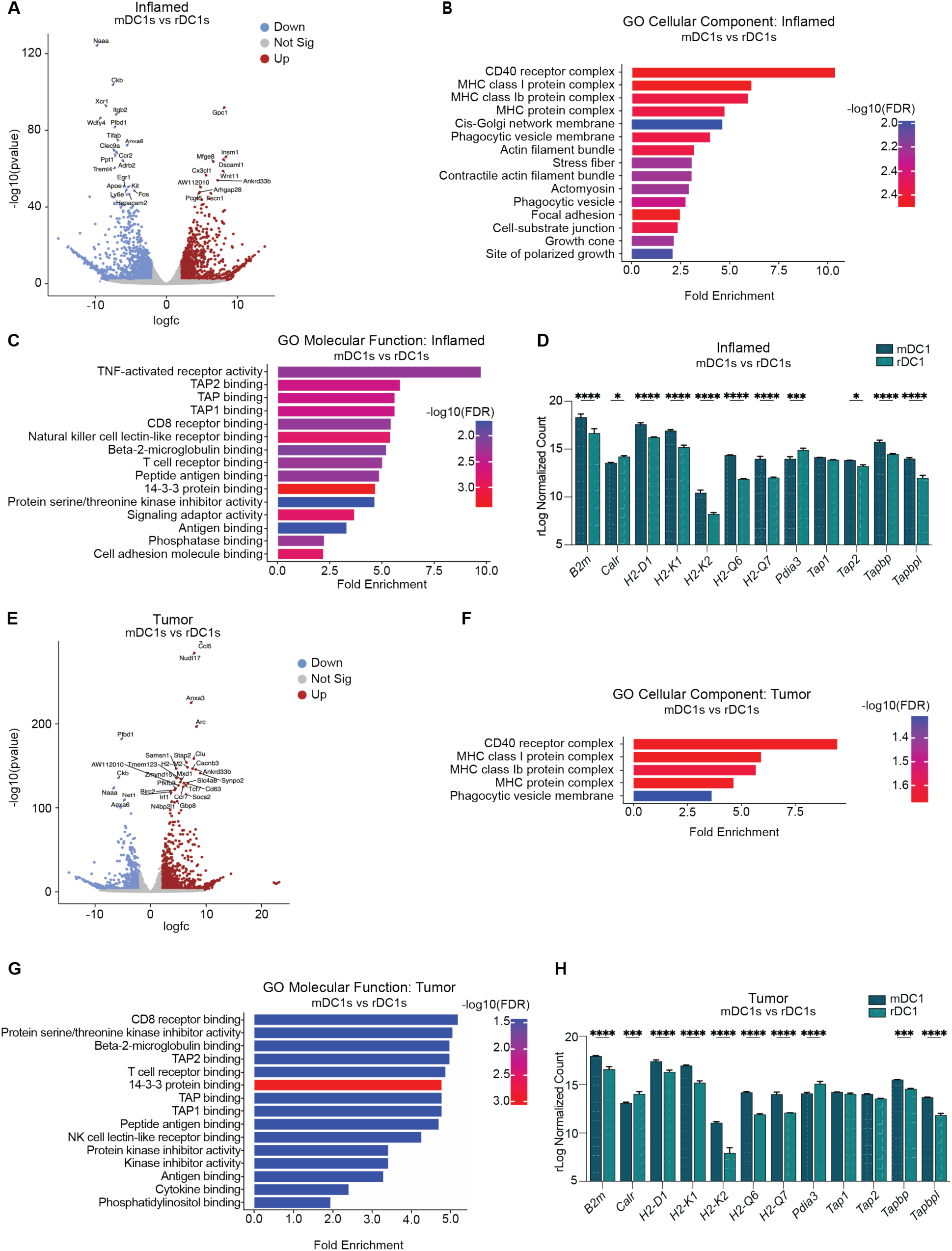
Type I mDCs display superior upregulation of peptide loading and presentation pathways compared to rDC. (**A**) Volcano plot of differentially expressed genes from mDC1s compared to rDC1s from dLNs of inflamed tissue microenvironments. (**B**) Pathway analysis of Cellular Components and (**C**) Molecular Function upregulated in mDC1s vs rDC1s from dLNs of inflamed tissues. (**D**) Normalized counts of antigen presentation genes from mDC1s and rDC1s from dLNs of inflamed tissues. (**E**) Volcano plot of differentially expressed genes from mDC1s compared to rDC1s from tumor dLNs. (**F**) Pathway analysis of Cellular Components and (**G**) Molecular Function upregulated in mDC1s vs rDC1s from dLNs of tumor tissues. (**H**) Normalized counts of antigen presentation genes from mDC1s and rDC1s from dLNs of tumor tissues. (A - C, E - G) Data are plotted as mean or (D, H) mean ± SD. n = 2 technical replicates; each replicate represents 20 mice (6 lymph nodes each) per tissue condition. (D, H) *p < 0.05, **p < 0.01, ***p < 0.001, ****p < 0.0001 by ordinary two-way ANOVA with Šídák multiple comparisons test. Migratory type 2 cDC, mDC2. Resident type 1 cDC, rDC1. Resident type 2 cDC, rDC2. ZsGreen, ZsG.

Given the large degree of antigen transfer from migratory to resident cDCs in tumor dLNs (Fig. 1B, 1G), determining differences between mDC1s and rDC1s is important for understanding how this antigen transfer impacts antitumor T cell responses. Thus, we next compared corresponding mDC1s and rDC1s derived from tumor dLNs (Fig. 4E). Similar analysis of these cellular components showed mDC1s were enriched in MHCI protein components and the costimulatory CD40 receptor complex compared to rDC1s (Fig. 4F). Top pathways in tumor dLN mDC1s were similar to those in the inflamed dLN mDC1s, primarily involved in peptide-MHCI loading and binding (Fig. 4G). As similar pathways and components were upregulated in mDC1s compared to rDC1s independent of inflamed or tumor tissue, we investigated the context-independent overlap between the cDC1 subsets by examining upregulated genes in mDC1s from both tissue states (Fig. S4A). Top cellular components and pathways in both peripheral tissue environments supported the conclusion that MHCI machinery and binding are key differences between mDC1s and rDC1s (Fig. S4B, S4C). These components and pathways have the potential to promote T cell proliferation through MHCI loading and presentation of antigen. Importantly, these cDC subset differences in gene expression regulating MHCI and peptide MHCI loading, alongside differences in surface MHCI proteins (Fig. 3J-3N), suggest that rDC1s fail to achieve the same antitumor T cell priming potential as mDC1s.

In summary, migratory cDCs, particularly mDC1s, are transcriptionally and functionally superior at cross-presenting skin tumor antigens compared to steady state or inflamed skin antigens. Resident cDCs, including rDC1s, fail to upregulate MHCI antigen presentation pathways in tumor dLNs to the same extent as mDC1s. Further, rDC1s exhibit lower baseline expression of MHCI machinery than mDC1s, which corresponds to a reduced per-cell capacity to prime naïve CD8^+^ T cells. These findings underscore the importance of harnessing the mDC1 subset for therapy and designing strategies to optimize the cell-intrinsic cross-presentation programs of additional cDC subsets. These data support the development of immunotherapies that enhance resident cDC cross-presentation efficiency of tumor antigen or selectively target antigen vaccines to mDC1 subsets in an effort to boost antitumor immunity.

## DISCUSSION

Tumor dLNs exhibit a qualitatively distinct pattern of antigen dispersal compared with steady state or acutely inflamed skin dLNs: tumor-derived antigen becomes broadly distributed across both migratory and resident cDC subsets, whereas steady state and DBP-induced inflammation result in antigen being concentrated primarily in migratory cDCs. We find that migratory cDC1s are transcriptionally and functionally optimized for cross-presentation, as evidenced by upregulated expression of MHCI machinery, peptide-loading components, CD40, and inflammatory cytokines including IL12. Further, migratory cDC1s drive the most potent CD8^+^ T cell proliferation. By contrast, resident cDCs in tumor dLNs receive substantial antigen but do not engage cross-presentation and DC maturation gene expression programs to the same degree nor do they elicit equivalent naïve T cell priming as measured by proliferation. Notably, our findings do not support an intrinsic deficiency in cross-presentation by resident cDC1s. When provided antigen *ex vivo*, rDC1s efficiently stimulated OTI proliferation and, unexpectedly, were the most potent subset at driving proliferation following uptake of whole OVA protein. These findings suggest that the diminished priming capacity of antigen-bearing resident cDCs observed *in vivo* is more likely attributable to differences in the form of antigen available to these cells or the distinct maturation state of the cDCs *in vivo* in a given context rather than an inherent inability to cross-present.

Previous studies have reported that both migratory and resident cDCs can present antigens and contribute to T cell responses, but the relative importance of each subset is context dependent^1,9,13^. Our data separate two interdependent determinants of priming efficacy: (1) which cDCs hold antigen in the dLN and (2) which cDCs are intrinsically competent to cross-present to CD8^+^ T cells. Tumors increase overall antigen availability in the dLN, but cDCs redistribute that antigen away from the subset best equipped to convert antigen into robust MHCI presentation and costimulation. The transcriptional signature of tumor-associated mDC1s provides a molecular explanation for their superior ability to drive T cell proliferation. Thus, antigen transfer in tumor dLNs may act as a double-edged sword, increasing antigen availability while delivering antigen to less competent presenters. Although proliferative capacity was most strongly associated with migratory cDC1s, we observed context-dependent differences in T cell phenotypes following priming by resident cDC subsets. These effects were modest and not uniformly maintained across comparisons, but they raise the possibility that resident cDCs contribute to qualitatively distinct CD8^+^ T cell differentiation programs. Indeed previous studies suggest that CD8^+^ T cell exhibit distinct transcriptional programs directly related to the subset of antigen-presenting cDC^10^. While technically challenging, future studies that deteriming cDC subset priming dynamics and downstream T cell function *in vivo* will be needed to fully disentangle the impact of antigen dispersal on the antitumor T cell responses.

An important unanswered question is how immunomodulatory therapies that directly target dendritic cells influence these antigen distribution patterns. In particular, CD40 agonists have been shown to enhance dendritic cell activation, licensing, and T cell priming, and our transcriptional analyses identified enrichment of CD40-associated pathways within migratory cDC1s. It will therefore be important in future studies to determine whether CD40 agonism alters the extent and functional consequences of antigen dispersal among cDC subsets within tumor dLNs.

Our findings show that the pattern of antigen dispersal is a determinant of downstream T cell priming. Antigen in mDC1s favors robust MHCI presentation and rapid, multiple CD8^+^ T cell divisions, whereas broad dispersal in tumor dLNs reallocates antigen to resident cDCs that lack the same cross-presentation program and consequently reduce per-cell priming efficiency. By uncoupling antigen availability from antigen cross-presentation competence, our study identifies LN antigen dispersal as a previously underappreciated mechanism that can impact antitumor CD8^+^ T cell priming. Strategies that preserve mDC1 antigen access or that enhance resident cDC cross-presentation programs offer complementary paths to potentially increase the magnitude and quality of antitumor T cell responses.

## Limitations of the study

While previous studies have demonstrated migratory-to-resident cDC antigen transfer in the B16-ZsGreen model, the current study was not designed to directly measure antigen transfer. Our analyses used several timepoints during inflammation and tumor growth, however, pulse-chase approaches and direct measurements of transfer kinetics are needed in order to fully resolve the dynamics of antigen transfer and persistence within individual cDC subsets. Additionally, whole LN imaging would be needed to better characterize the role of spatial localization in driving antigen transfer and establishing cDC phenotypes and cDC:T cell interactions. Further, functional readouts focused on *ex vivo* coculture and bulk transcriptional profiling; definitive causal links between antigen dispersal and *in vivo* tumor control will require subset-restricted antigen delivery or blockade of antigen transfer combined with tumor outcome measures.

Our study examined two transplantable tumor models and one model of acute skin inflammation. Although the broad pattern of antigen dispersal was conserved across B16-ZsGreen and MB49-ZsGreen tumors, future studies using genetically engineered mouse models and tumors spanning immune “hot” and “cold” states will be important to determine the generalizability of these findings and the connection with the downstream adaptive immune response. In addition, steady state and inflamed conditions were studied in K14-Cre; Rosa26-ZsGreen mice, whereas tumor studies were performed in wildtype recipients bearing fluorescent tumors. Although we are unaware of evidence that the Rosa26-ZsGreen allele alters dendritic cell biology, we cannot formally exclude a contribution of host genotype in this setting.

Our studies used ZsGreen and OVA as reporter antigens to enable tracking of antigen acquisition and antigen-specific CD8^+^ T cell responses. While previous work has shown that ZsGreen co-localizes with known endogenous tumor antigens in cDCs^6^, tumor neoantigens may differ in abundance, stability, or processing and could show different dispersal or presentation dynamics. Furthermore, our analyses focused primarily on MHCI-restricted antigen presentation and CD8^+^ T cell priming; therefore, the impact of antigen dispersal on MHCII presentation and CD4^+^ T cell responses remains unresolved. Finally, our studies do not address human cDCs and thus future studies using human cells will be needed to establish translational relevance.

## RESOURCE AVAILABILITY

### Lead contact

Requests for further information and resources should be directed to and will be fulfilled by the lead contact, Megan K Ruhland.

### Materials availability

All unique/stable reagents generated in this study are available from the lead contact upon request.

### Data and code availability

- RNA-seq data have been deposited at the NCBI Sequence Read Archive (SRA) as accession number: PRJNA1498663. Data are publicly available as of the date of publication.
- This paper does not report original code.
- Any additional information required to reanalyze the data reported in this paper is available from the lead contact upon request.

## ACKNOWLEDGMENTS

We thank M. Krummel for providing the K14-ZsGreen mice and B16F10 reporter cell lines. We thank E. Alspach for discussion and support. Funding: This work was supported by Harry J. Lloyd Charitable Trust (M.K.R.), the V Foundation V Scholar Award (M.K.R), the Concern Foundation Conquer Cancer Now Award (M.K.R.), NCI R01CA299335 (M.K.R.), T32AI170496 (V.P.S.) and T32GM142619 (K.B.). We acknowledge the OHSU Flow Cytometry Shared Resource (RRID SCR_009974) for assistance generating flow cytometry data and the Massively Parallel Sequencing Shared Resource (RRID SCR_009984) for generating the RNA sequencing data. Research reported here was supported in part by the Knight Cancer Institute CCSG grant NIH P30CA069533 and NIH grant S10OD028512.

## AUTHOR CONTRIBUTIONS

Conceptualization, M.K.R.; methodology, V.P.S and M.K.R; Investigation, V.P.S, N.B., K.B., C.R., V.L., J.R.U. and M.K.R; writing – original draft, V.P.S and M.K.R; writing – review & editing; V.P.S, N.B. and M.K.R; funding acquisition, V.P.S., K.B. and M.K.R.; supervision, M.K.R.

## SUPPLEMENTAL INFORMATION

Document S1. Supplemental Figures S1-S4

## METHODS

KEY RESOURCES TABLE.

| REAGENT or RESOURCE | SOURCE | IDENTIFIER |
| --- | --- | --- |
| **Antibodies** |  |  |
| Biotin anti-mouse CD2 Antibody | BioLegend | Cat#100104; RRID:AB_312651 |
| BD Pharmingen™ Purified Rat Anti-Mouse CD16/CD32 (Mouse BD Fc Block™) | BD Biosciences | Cat#553142; RRID:AB_394657 |
| PerCP/Cyanine5.5 anti-mouse CD8a Antibody | BioLegend | Cat#100734; RRID:AB_2075238 |
| Alexa Fluor® 700 anti-mouse I-A/I-E Antibody | BioLegend | Cat#107622; RRID:AB_493727 |
| Brilliant Violet 421™ anti-mouse H-2K<sup>b</sup> Antibody | BioLegend | Cat#116525; RRID:AB_2876430 |
| Brilliant Violet 605™ anti-mouse/human CD11b Antibody | BioLegend | Cat#101257; RRID:AB_2565431 |
| Brilliant Violet 650™ anti-mouse CD11c Antibody | BioLegend | Cat#117339; RRID:AB_2562414 |
| Brilliant Violet 711™ anti-mouse Ly-6C Antibody | BioLegend | Cat#128037; RRID:AB_2562630 |
| APC anti-mouse CD103 Antibody | BioLegend | Cat#121414; RRID:AB_1227502 |
| PE/Cyanine7 anti-mouse CD45 Antibody | BioLegend | Cat#103114; RRID:AB_312979 |
| BD Horizon™ BUV395 Rat Anti-Mouse CD86 | BD Biosciences | Cat#564199; RRID:AB_2738664 |
| Brilliant Violet 785™ anti-mouse/human CD45R/B220 Antibody | BioLegend | Cat#103246; RRID:AB_2563256 |
| Brilliant Violet 785™ anti-mouse Ly-6G Antibody | BioLegend | Cat#127645; RRID:AB_2566317 |
| Brilliant Violet 785™ anti-mouse NK-1.1 Antibody | BioLegend | Cat#108749; RRID:AB_2564304 |
| Brilliant Violet 785™ anti-mouse CD90.2 (Thy1.2) Antibody | BioLegend | Cat#105331; RRID:AB_2562900 |
| BD Pharmingen™ APC Rat Anti-Mouse IL-12 (p40/p70) | BD Biosciences | Cat#554480; RRID:AB_398560 |
| IL-1 beta (Pro-form) Monoclonal Antibody (NJTEN3), PE, eBioscience™ | Thermo Fisher Scientific | Cat#12-7114-82; RRID:AB_10732630 |
| Brilliant Violet 711™ anti-mouse CD103 Antibody | BioLegend | Cat#121435; RRID:AB_2686970 |
| BD Horizon™ BUV737 Rat Anti-Mouse CD44 | BD Biosciences | Cat#612799; RRID:AB_2870126 |
| Brilliant Violet 711™ anti-mouse CD8a Antibody | BioLegend | Cat#100759; RRID:AB_2563510 |
| Brilliant Violet 605™ anti-mouse CD279 (PD-1) Antibody | BioLegend | Cat#135220; RRID:AB_2562616 |
| BD Horizon™ BUV395 Mouse Anti-EOMES | BD Biosciences | Cat#567171; RRID:AB_2916488 |
| PerCP/Cyanine5.5 anti-T-bet Antibody | BioLegend | Cat#644806; RRID:AB_1595488 |
| Brilliant Violet 421™ anti-mouse IFN-γ Antibody | BioLegend | Cat#505830; RRID:AB_2563105 |
| FITC anti-mouse TNF-α Antibody | BioLegend | Cat#506304; RRID:AB_315425 |
| Granzyme B Monoclonal Antibody (NGZB), APC, eBioscience™ | Thermo Fisher Scientific | Cat#17-8898-82; RRID:AB_2688068 |
| **Biological samples** |  |  |
| Mouse tissue samples (LN, spleen) | OHSU | N/A |
| **Chemicals, peptides, and recombinant proteins** |  |  |
| dibutyl-phthalate (DBP) | Sigma Aldrich | Cat#NC1014483 |
| DMEM | Gibco | Cat#11995065 |
| Fetal bovine serum | HyClone | Cat#SH3039603 |
| Penicillin-Streptomycin-Glutamine | Gibco | Cat#10378-016 |
| RPMI1640 | Gibco | Cat#11875-119 |
| Matrigel matrix | Corning | Cat#356231 |
| Collagenase, Type I | Worthington Biochemical | Cat#LS004197 |
| Collagenase, Type IV | Worthington Biochemical | Cat#LS004189 |
| DNAse I | Roche | Cat#10104159001 |
| EDTA | Fisher BioReagents | Cat#BP2482100 |
| HEPES | MilliporeSigma | Cat#H0887-100ML |
| 2-mercaptoethanol | Gibco | Cat#21985023 |
| Ovalbumin | Worthington Biochemical | Cat#LS003061 |
| OVA (257-264) Peptide Fragment | Anaspec | Cat#AS-60193-1 |
| Lipopolysaccharides from Escherichia coli | Sigma-Aldrich | Cat#L6529-1MG |
| ACK lysing buffer | Gibco | Cat#A1049201 |
| Critical commercial assays |  |  |
| Zombie NIR Fixable Viability Kit | BioLegend | Cat#423105 |
| CD8<sup>+</sup> T cell isolation kit | STEMCELL Technologies | Cat#19853 |
| QIAshredder | Qiagen | Cat#79654 |
| RNeasy Micro Kit | Qiagen | Cat#74004 |
| Violet Proliferation Dye | BD Biosciences | Cat#562158 |
| EasySep Mouse Streptavidin RapidSpheres Isolation Kit | STEMCELL Technologies | Cat#19860 |
| Cytofix/Cytoperm Solution | BD Biosciences | Cat#555028 |
| Cytofix Fixation Buffer | BD Biosciences | Cat#554655 |
| BD GolgiPlug™ Protein Transport Inhibitor (Containing Brefeldin A) | BD Biosciences | Cat#555029 |
| Deposited data |  |  |
| RNA-sequencing dataset | This paper | BioProject: PRJNA1498663 |
| Experimental models: Cell lines |  |  |
| B16-F10 | ATCC | Cat#CRL-6475; RRID:CVCL_0159 |
| B16-ZsGreen | OHSU | N/A |
| B16-ZsGreen-OVA | OHSU | N/A |
| B16-mCherry | OHSU | N/A |
| Experimental models: Organisms/strains |  |  |
| Mouse: C57BL/6 | The Jackson Laboratory | RRID:IMSR_JAX:000664 |
| Mouse: OTI (C57BL/6-Tg(TcraTcrb)1100Mjb/J | The Jackson Laboratory | RRID:IMSR_JAX:003831 |
| Mouse: K14-Cre; lox-stop-lox-ZsGreen | M. Krummel<sup>10</sup> | N/A |
| Mouse: C57BL/6-Tg(KRT14-OVAL*)15Sika/J | The Jackson Laboratory | RRID:IMSR_JAX:026562 |
| Software and algorithms |  |  |
| FlowJo, v.10 | Becton Dickinson | https://www.flowjo.com/ |
| Prism, v.10 | GraphPad | https://www.graphpad.com/features |
| Galaxy | N/A | usegalaxy.org |
| ShinyGo, v0.82 | N/A | https://bioinformatics.sdstate.edu/go/ |

## EXPERIMENTAL MODEL DETAILS

### Mice

Mice were housed in a specific pathogen-free environment at the Oregon Health and Science University (OHSU) mouse facility and all experimentation methods were approved by the The Institutional Animal Care and Use Committee (IACUC) at OHSU. Mice used for the study include: C57BL/6 (bred in house or purchased from Jackson Laboratory, Strain #000664), OTI (C57BL/6-Tg(TcraTcrb)1100Mjb/J, bred in house or purchased from Jackson Laboratory, Strain #003831), K14-Cre; lox-stop-lox-ZsGreen mice which express cytoplasmic ZsGreen in a keratinocyte restricted manner; animals were a kind gift from M. Krummel ^10^, and K14-Cre; lox-stop-lox-ZsGreen; K14-OVA mice were generated by crossing K14-ZsGreen animals with C57BL/6-Tg(KRT14-OVAL*)15Sika/J from Jackson Labs, Strain #026562 (K14-OVA). A combination of 6-12 week old male and female mice were used for experiments.

### Cell lines

B16-ZsGreen and B16-ZsGreen-OVA cell lines were generated and validated as previously described ^10^. Briefly, the B16-F10 cell line (ATCC, CRL-6475) was virally transduced with constructs for ZsGreen or ZsGreen-OVA, respectively, and FACS sorted to select for stable genetic expression. Cell lines were housed in incubators at 37°C, 5% CO_2_, and 95% humidity. Cell lines were maintained with D10 media: DMEM (Gibco, 11995065), 10% heat-inactivated FBS (HyClone, SH3039603), and 1% Penicillin-Streptomycin-Glutamine (Gibco, 10378-016). MB49-ZsGreen cell lines were generated similarly and maintained with D10 supplemented with 1% each of HEPES solution (MilliporeSigma, H0887-100ML), minimum essential media (MEM) non-essential amino acids solution (Sigma-Aldrich, S8636-100ML), and sodium pyruvate solution (Sigma-Aldrich, S8636-100ML).

## METHODS DETAILS

### Tumor injections

B16-ZsGreen, MB49-ZsGreen or B16-ZsGreen-OVA lines were harvested at confluency, washed with PBS twice and resuspended to 10^7^ cells/mL in RPMI1640 (Gibco, 11875-119). Cells were diluted with Matrigel matrix (Corning, 356231) for a 1:1 mixture. Mixture was used to perform bilateral injections subcutaneously in both flanks of C57BL/6 mice at 50 μL, 2.5×10^5^ cells per injection. B16-ZsGreen and MB49-ZsGreen injections were grown for 14 days and B16-ZsGreen-OVA injections were grown for 21 days prior to harvest of skin dLNs.

### Topical applications

Dorsal skin of K14-ZsGreen or K14-ZsGreen-OVA mice were shaved one day prior to beginning of topical applications. A 1:1 mixture of dibutyl-phthalate, DBP (Sigma Aldrich, NC1014483) and acetone was made, and 200 μL of the DBP mixture or acetone vehicle control was applied dropwise to the shaved backs of the mice. Topical applications were performed daily for a total of 7 days prior to harvest of skin dLNs.

### Lymph node digest

Inguinal, axillary, and brachial LNs were harvested from tumor bearing C57BL/6 mice, treated K14-ZsGreen, or treated K14-ZsGreen-OVA mice. LN harvests and digests were performed as previously described ^6^. Briefly, for each mouse, LNs were harvested and placed into one well of a 24 well plate (Corning, 3524) containing 1mL RPMI1640 (Gibco, 11875-119) on ice. For each mouse, 1mL digest buffer was prepped in RPMI1640: 100U Collagenase, Type I (Worthington Biochemical, LS004197), 500U Collagenase, Type IV (Worthington Biochemical, LS004189), and 20 μg DNAse I (Roche, 10104159001). Cleaned LNs were punctured with sharp forceps to open capsule and placed into 1mL digest buffer in a well of the 24 well plate. Plates were incubated at 37°C for 30 minutes, pipetting up and down each well halfway through incubation time. LNs were then washed and filtered through a 70 μm cell strainer (Corning 431751) with MACs buffer: PBS (Gibco, 14190144) supplemented with 2% heat-inactivated FBS (HyClone, SH3039603), 1mM EDTA (Fisher BioReagents, BP2482100), 1% Penicillin-Streptomycin-Glutamine (Gibco, 10378-016), and 2mM HEPES solution (MilliporeSigma, H0887-100ML). Cells for flow cytometry were then stained in MACs buffer. Cells for FACS were negatively selected for CD2 using a biotin-conjugated anti-CD2 antibody (BioLegend, 100104) alongside an EasySep Mouse Streptavidin RapidSpheres Isolation Kit (STEMCELL Technologies, 19860) and magnetic depletion (STEMCELL Technologies, 18000), per manufacturer’s protocol, and then stained.

### Cell staining

For intracellular staining, cells in a 96 well v-bottom plate (Corning, 3894) were spun down and resuspended in R10: RPMI1640 (Gibco, 11875-119), 10% heat-inactivated FBS (HyClone, SH3039603), 1% Penicillin-Streptomycin-Glutamine (Gibco, 10378-016), 0.1% 2-mercaptoethanol (Gibco, 21985023). GolgiPlug (BD Biosciences, 555029) was added to the resuspension media at 1 μL/mL and returned to incubator for 5 hours. Cells were then spun down and incubated for 15 minutes on ice with viability dye (BioLegend, 423105). Cells were then stained for surface markes for 20 minutes on ice with Fc Block (BD Biosciences, 553142) followed by permeabilization (if applicable) by resuspending cells in Cytofix/Cytoperm Solution (BD Biosciences, 554655) for 20 minutes on ice. Permeabilized cells were then stained with intracellular antibodies and Fc Block diluted in the Cytofix/Cytoperm Wash Buffer for 1 hour on ice. All samples were washed and resuspended in MACs buffer before acquisition with flow cytometry or cell sorting.

### Flow cytometry and sorting

Cell sorting was performed with the BD FACSymphony S6 (BD Biosciences) with the 100 μm size nozzle and BD FACSDiva Software (BD Biosciences, v.9). Flow cytometry was performed with the BD FACSymphony A5 (BD Biosciences). FlowJo Software (Treestar, v.10) was used for flow cytometry analysis and proliferation modeling.

### T cell stimulation assays

Spleens and skin dLNs were harvested from OTI mice. Tissues were mashed with syringe plunger. Splenic tissues were incubated in ACK lysing buffer (Gibco, A1049201) briefly and all samples were brought up to 10 mL with MACs buffer. Cells were pelleted and filtered through a 70 μm cell strainer (Corning, 431751) and CD8^+^ T Cells were isolated using a CD8^+^ T cell isolation kit (STEMCELL Techologies, 19853). Isolated T cells were incubated with 1 μM Violet Proliferation Dye (BD Biosciences, 562158) in PBS for 15 minutes at for at 37°C. Cells were resuspended to 5×10^5^ cells/mL in R10. When indicated, sorted cDCs were cultured in D10 containing 8μg/mL of whole ovalbumin protein for 4 hours (Worthington Biochemical, LS003061) or 100 ng/mL OVA (257-264) peptide (SIINFEKL) for 1 hour (Anaspec, AS-60193-1) prior to washing with MACs buffer. Sorted cDCs were washed and resuspended to 5×10^4^ cells/mL in R10. T cell stimulation assays were set up at a 1:10 ratio in v-bottom plates with 100μL from each cell suspension (200μL total per well). Plates were incubated at 37°C, 5% CO_2_, and 95% humidity for 3 days prior to harvest. T cell activation was based off the gating strategy outlined in Figure 2A.

### Lymph Node Stimulation Assay

Inguinal, axillary, and brachial LNs from wildtype C57BL/6 mice were harvested and digested as described, and resuspended to 200μL D10 per bilateral set of LNs: DMEM (Gibco, 11995065), 10% heat-inactivated FBS (HyClone, SH3039603), 1% Penicillin-Streptomycin-Glutamine (Gibco, 10378-016), 0.1% 2-mercaptoethanol (Gibco, 21985023). Resuspended LNs were then transferred to 96-well plates and supplemented with 1μg/mL lipopolysaccharide (LPS) (Sigma-

Aldrich, L6529) for stimulation. Digested LNs were incubated at 37°C, 5% CO_2_, and 95% humidity for 5 hours with LPS prior to washing with MACs buffer and staining for flow cytometry analysis of cDC subsets and maturation markers.

### RNA-sequencing

ZsGreen^+^ cDC subsets were sorted as outlined in Figure 1A, and the QIAshredder (Qiagen, 79654) was used to homogenize samples after lysis. RNA was extracted from 1×10^4^ cell sorted cells with the RNeasy Micro Kit (Qiagen, 74004), per manufacturer’s protocol. Concentration and integrity measurement of each RNA sample was assessed using the Agilent Bioanalyzer 2100 with the Pico Chip assay. Library preparations and sequencing were performed by the Integrated Genomics Laboratory at Oregon Health and Science University with the SMARTSeq v4 PLUS Low Input RNA Kit (Takara) and the Illumina NextSeq 500 with 50 million read pairs per library. Sequencing files were uploaded to the Galaxy online platform at usegalaxy.org for processing ^29^. Within the Galaxy platform, Trimmomatic was used to trim adapter sequences of paired-end reads, HISAT2 was used to align reads to the mouse reference genome (GRCm38, mm10), htseq-count was used to count aligned reads in a BAM file, and DESeq2 was used for differential gene expression. The enrichment tool ShinyGo, v.0.82 (https://bioinformatics.sdstate.edu/go/) was used for Gene Ontology enrichment analysis.

## QUANTIFICATION AND STATISTICAL ANALYSIS

Statistical tests and data visualization were performed using GraphPad Prism (v. 10). All statistical analyses and descriptions of displayed data are declared within the figure legends. Data are presented as mean ± SEM, unless otherwise indicated. ANOVA and Student’s *t* tests with appropriate indicated corrections were used for comparison between groups to determine statistical significance. Biological replicates refer to individual mice. Significance was defined in the following manner: not significant (ns), *p < 0.05, **p < 0.01, ***p < 0.001, ****p < 0.0001, as determined by the appropriate statistical test.

