## Supplementary material for "Contextual control of CD8^+^ T cell priming by dendritic cell subsets in tumor and inflammatory microenvironments": Document S1. Supplemental Figures S1-S4

### Figure S1

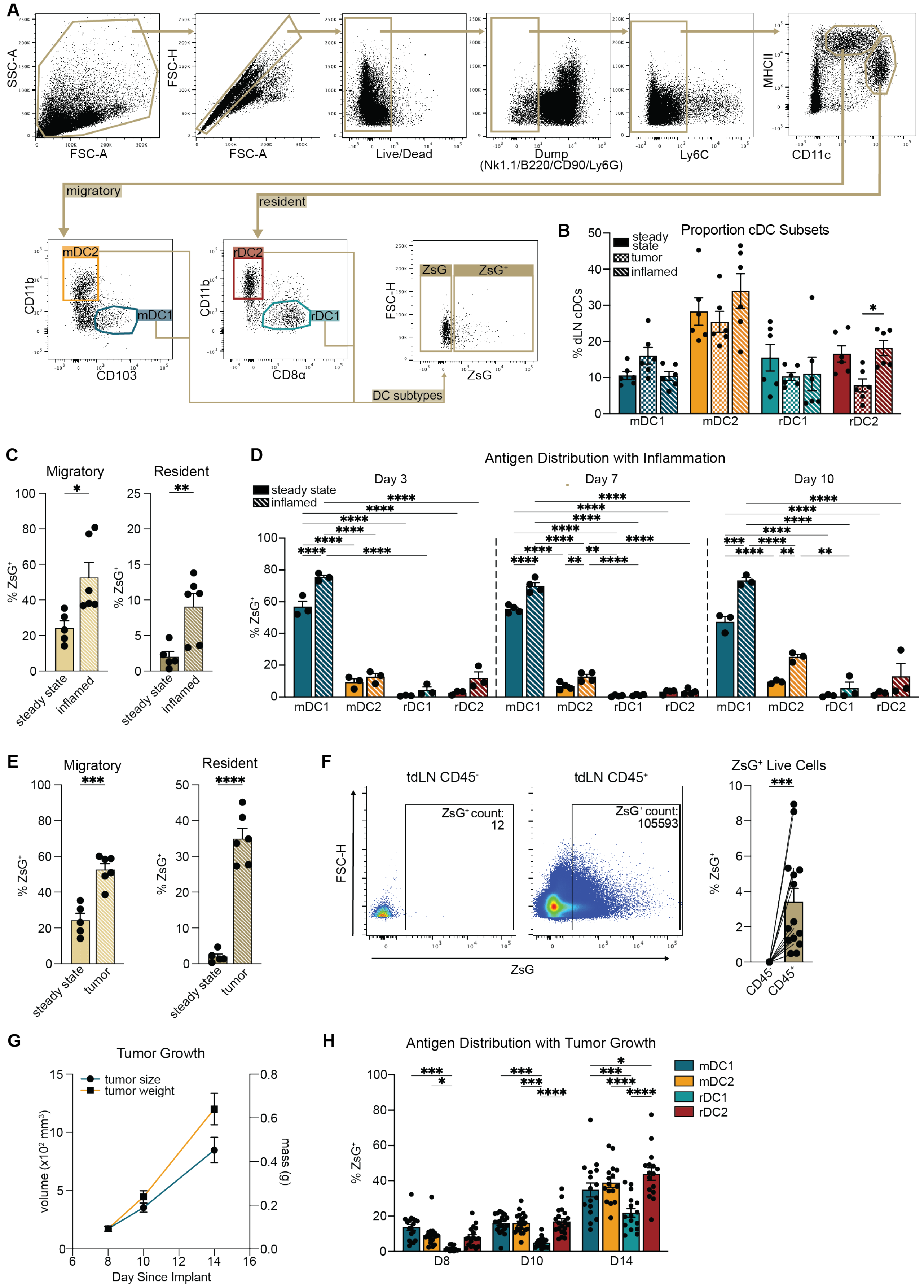

**Figure S1. Related to Figure 1.** (A) Gating strategy for ZsGreen antigen acquisition by Live/Nk1.1$^-$/B220$^-$/CD90$^-$/Ly6G$^-$/Ly6C$^-$ cDC subsets from draining lymph nodes (dLNs). (B) Flow cytometry analysis showing percent cDC subset composition from dLN of steady state, B16-ZsGreen tumor, and inflamed tissue contexts. (C) Percent ZsGreen$^+$ migratory (left) or resident (right) cDC subsets harvested from dLNs of inflamed or steady state skin tissue. (D) Percent ZsGreen$^+$ cDC subsets from skin dLNs 3, 7, or 10 days post initiation of topically induced inflammation. (E) Percent ZsGreen$^+$ migratory (left) or resident (right) cDC subsets harvested from dLNs of tumor or steady state skin tissue. (F) Representative flow cytometry plots (left) and quantification of CD45$^-$ and CD45$^+$ cells positive for ZsGreen (right), displayed as percent of live cells within dLNs, 14 days following B16-ZsGreen tumor implantation. (G) Tumor size and weight alongside (H) corresponding percent ZsGreen$^+$ of each cDC subset within tumor dLNs 8, 10, or 14 days post-implantation of B16-ZsGreen. (B - H) Data are plotted as mean ± SEM. (B - C, E) n= 5 (steady state) or n = 6 (inflamed or B16 tumor) mice per tissue condition, pooled data from 2 independent experiments. (D) Representative experiment shown, n = 3-4 mice per steady state or inflamed tissue condition per timepoint. 2 independent experiments. (F - H) (F) n = 7 mice or (G - H) n = 9 mice for each timepoint, pooled data from 2 independent experiments. *p < 0.05, **p < 0.01, ***p < 0.001, ****p < 0.0001 by (B, D, H) ordinary two-way ANOVA with (B, H) Tukey's multiple comparisons test or (D) Šídák multiple comparisons test, or by (C, E) Gaussian unpaired or (F) paired two-tailed t test. Migratory type 2 cDC, mDC2. Resident type 1 cDC, rDC1. Resident type 2 cDC, rDC2. ZsGreen, ZsG.

### Figure S2

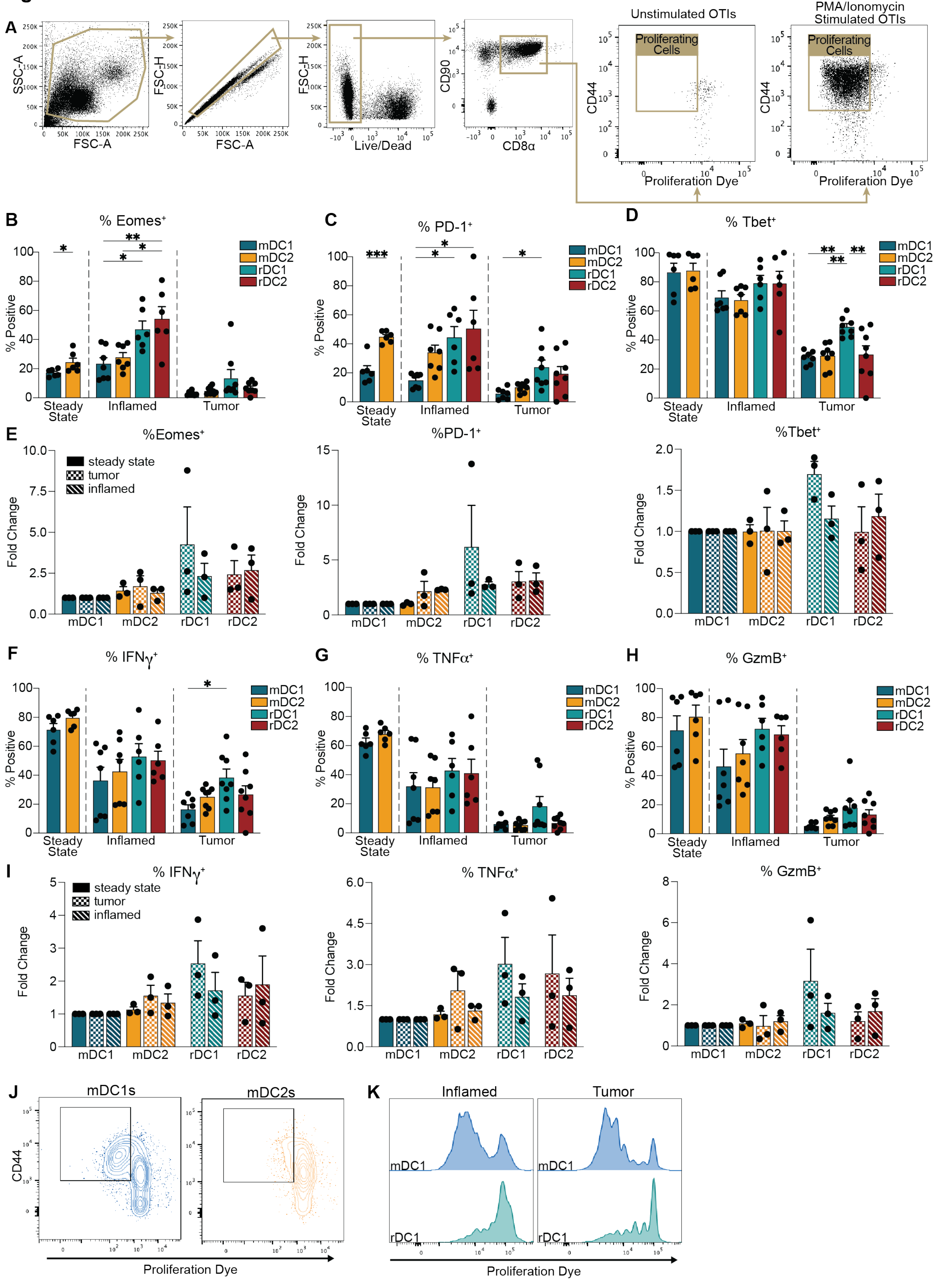
A
SSC-A
FSC-A
FSC-H
FSC-A
FSC-H
Live/Dead
CD90
CD8α
Unstimulated OTIs
PMA/Ionomycin Stimulated OTIs
Proliferating Cells
CD44
Proliferation Dye
B
% Eomes+
C
% PD-1+
D
% Tbet+
% Positive
mDC1
mDC2
rDC1
rDC2
Steady State
Inflamed
Tumor
E
%Eomes+
%PD-1+
%Tbet+
steady state
tumor
inflamed
Fold Change
F
% IFNγ+
G
% TNFα+
H
% GzmB+
I
% IFNγ+
% TNFα+
% GzmB+
J
mDC1s
mDC2s
K
Inflamed
Tumor
Proliferation Dye

**Figure S2. Related to Figure 2.** (A) Proliferation assay gating strategy for Live/$CD8^+$/$CD90^+$/$CD44^+$ proliferating OTI cells. (B - D) Flow cytometry quantification of percent proliferating $CD8^+$ T cells positive for (B) Eomes, (C) PD-1 and (D) Tbet, following coculture with sorted $ZsGreen^+$ cDCs. (E) Fold change for Eomes (left), PD-1 (middle) and Tbet (right) normalized to mean mDC1-induced proliferation in a given experiment under the indicated treatment. (F-H) Flow cytometry quantification of percent proliferating $CD8^+$ T cells positive for (F) interferon γ (IFNγ), (G) tumor necrosis factor α (TNFα), and (H) granzyme B (GzmB) following coculture with sorted $ZsGreen^+$ cDCs. (I) Fold change for IFNγ (left), TNFα (middle), and GzmB (right), normalized to mean mDC1-induced proliferation in a given experiment under the indicated treatment. (J) Representative flow cytometry plot for proliferation analysis of OTIs by cDC subsets sorted from mice under steady state conditions. (K) Representative proliferation histograms comparing OTI $CD8^+$ T cell cocultures with mDC1s and rDC1s sorted from inflamed (left) or B16-ZsGreen tumor (right) conditions. (B - I) Data are plotted as mean ± SEM. One to three technical replicates from cDCs sorted from 6 pooled lymph nodes of 10 mice per tissue condition for each experiment, n = 6 - 8 from pooled data from 3 independent experiments. (E, I) Data points represent the mean of each individual experiment. *$p < 0.05$, **$p < 0.01$, ***$p < 0.001$, ****$p < 0.0001$ by (B - D, F - H) individual ordinary one-way ANOVAs or (E, I) ordinary two-way ANOVA with (B - I) Tukey's multiple comparisons test. Migratory type 2 cDC, mDC2. Resident type 1 cDC, rDC1. Resident type 2 cDC, rDC2. ZsGreen, ZsG.

**Figure S3**

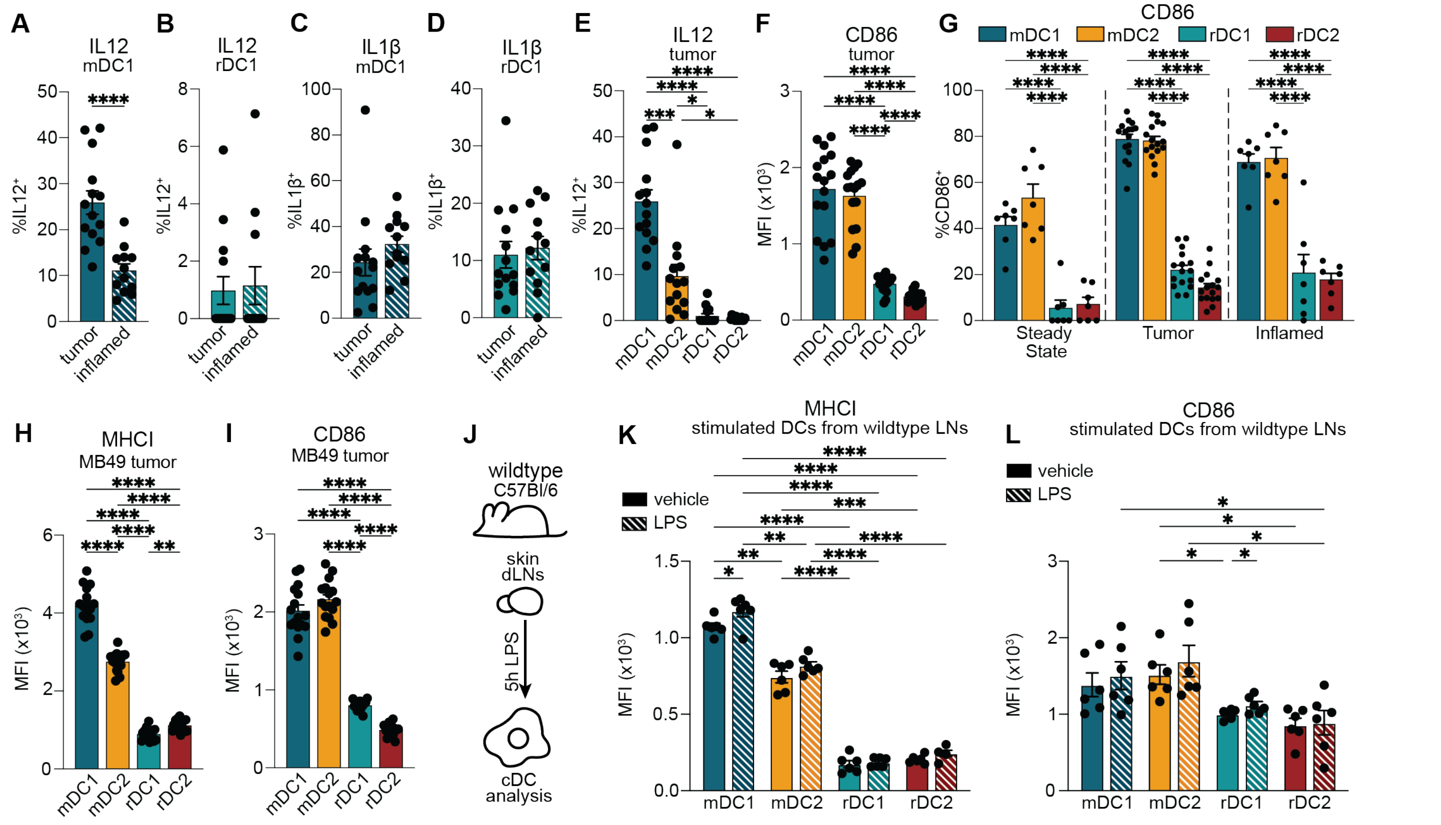


**Figure S3. Related to Figure 3.** (A-B) Percentage IL12+ (A) mDC1s and (B) rDC1s from dLNs of inflamed and B16-ZsGreen tumor conditions by flow cytometry. (C-D) Percentage IL1β+ (C) mDC1s and (D) rDC1s from dLNs of inflamed and B16-ZsGreen tumor conditions. (E) Percentage IL12+ cDCs by subset from B16-ZsGreen tumor dLNs by flow cytometry. (F) MFI of costimulatory molecule CD86 by cDC subset from tumor dLNs. (G) Percentage CD86+ of dLN cDCs from tumor, inflamed, and steady state conditions. (H - I) MFI of (H) MHCI and (I) CD86 for cDC subsets isolated from dLNs of MB49-ZsGreen tumors. (J) Schematic for *ex vivo* wildtype dLN stimulation with LPS and subsequent flow cytometry analysis. (K - L) Flow cytometry analysis for (K) MHCI and (L) CD86 MFI of cDC subsets from vehicle or LPS treated dLNs. (A - I, K - L) Data are plotted as mean ± SEM. (A - E, H - I) n = 6 (inflamed) or 8 (B16-ZsGreen or MB49-ZsGreen tumor) samples, data pooled from 2 independent experiments. (F - G) n = 7 (steady state or inflamed mice) or 8 (tumors) samples, data pooled from 2 independent experiments. (K - L) n = 6 biological replicates, data pooled from 2 independent experiments. *p < 0.05, **p < 0.01, ***p < 0.001, ****p < 0.0001 by (A - D) Welch's two-tailed t test, or (E - F, H - I) RM one-way ANOVA with Geisser-Greenhouse correction, (G) ordinary two-way ANOVA, or (K - L) RM two-way ANOVA with Geisser-Greenhouse correction with (E - I) Tukey's or (K - L) Šídák multiple comparisons test. Migratory type 2 cDC, mDC2. Resident type 1 cDC, rDC1. Resident type 2 cDC, rDC2. ZsGreen, ZsG.

### Figure S4

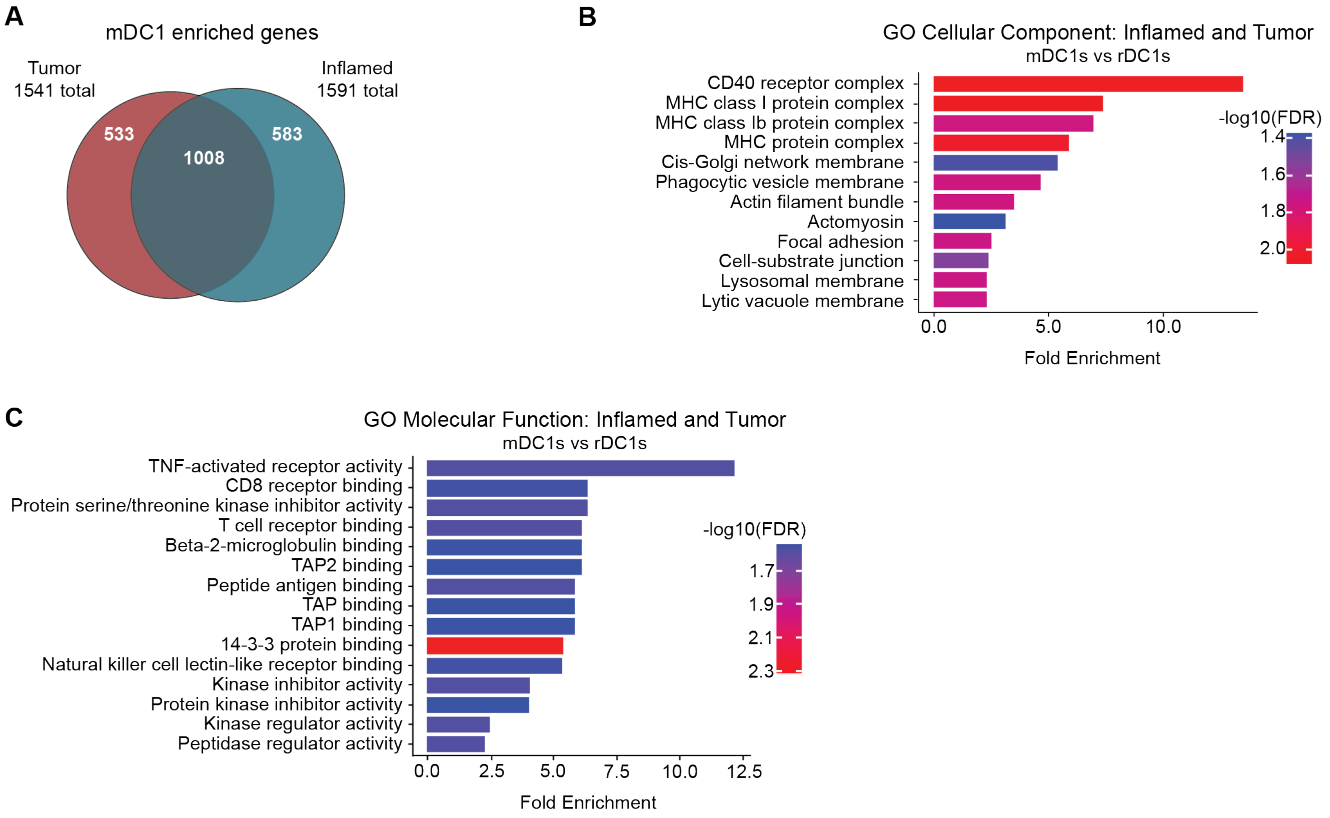


**Figure S4. Related to Figure 4.** (A) Venn diagram of upregulated genes as analyzed by DESeq2 in migratory cDC1s vs resident cDC1s from dLNs of tumor and inflamed conditions. (B) Pathway analysis of Cellular Components and (C) Molecular Function upregulated in mDC1s versus rDC1s from dLNs of both inflamed and tumor tissues. (A - C) Data are plotted as mean. n = 2 technical replicates; each replicate represents 20 mice (6 lymph nodes each) per tissue condition.
